## Supplementary material for "diaTracer enables spectrum-centric analysis of diaPASEF proteomics data": Description of Additional Supplementary Files

**Title:** Supplementary Data 1

**Description:** Proteins containing semi-tryptic peptides quantified by FP-diaTracer semi-tryptic workflow from the CSF dataset. The first column is the protein entry name. The second column is the number of semi-tryptic peptides quantified in the protein. The third column is the number of tryptic peptides quantified in the protein. The fourth column is the ratio between numbers of quantified semi-tryptic and tryptic peptides. The following columns are attributes of proteins from Uniprot.

**Title:** Supplementary Data 2

**Description:** Proteins containing semi-tryptic peptides mapping to the region immediately following the signal peptides in the N-terminal portion.

**Title:** Supplementary Data 3

**Description:** Protein differential expression results downloaded from FragPipe-Analyst for FP-diaTracer tryptic (Sheet 1) and FP-diaTracer semi-tryptic (Sheet 2) workflow.
