## Supplementary Figures for "diaTracer enables spectrum-centric analysis of diaPASEF proteomics data"

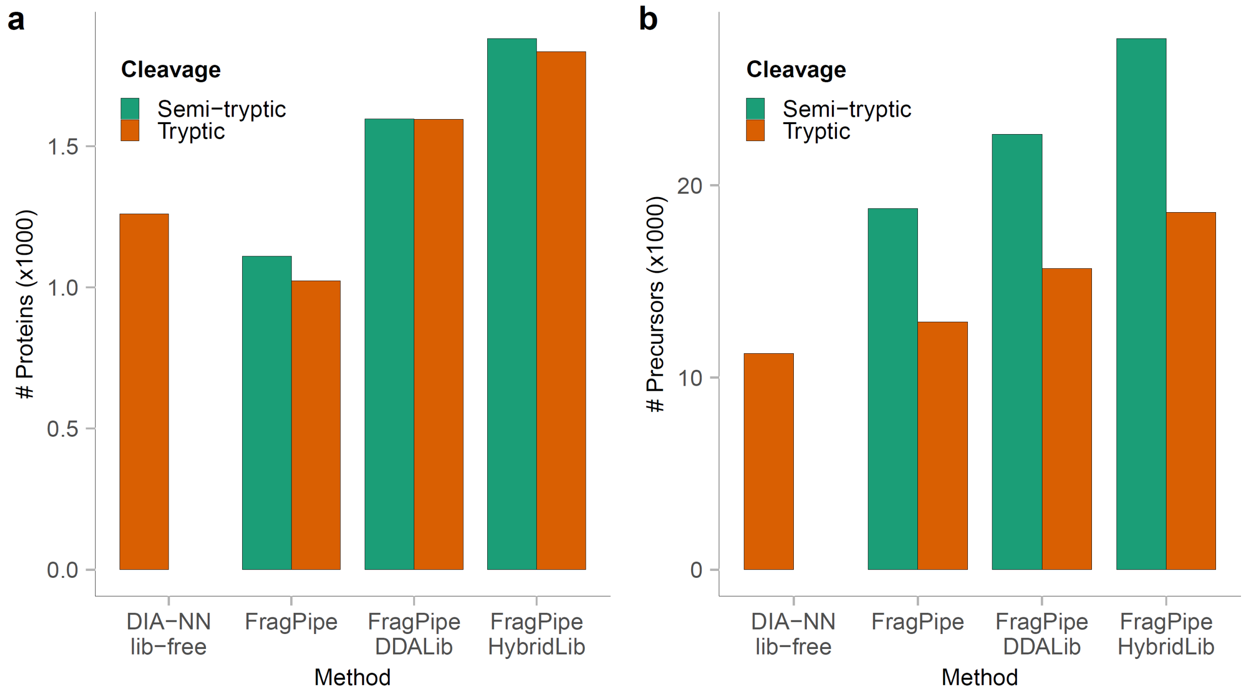


**Supplementary Figure 1.** Performance evaluation of tryptic, semi-tryptic, and mass offset search using a CSF dataset. **a)** Bar plot showing the total number of quantified proteins using different methods, with colors representing cleavage types (green: strict trypsin cleavage; orange: semi-tryptic cleavage). **b)** Bar plot showing the total number of precursors quantified in the CSF dataset using different methods.

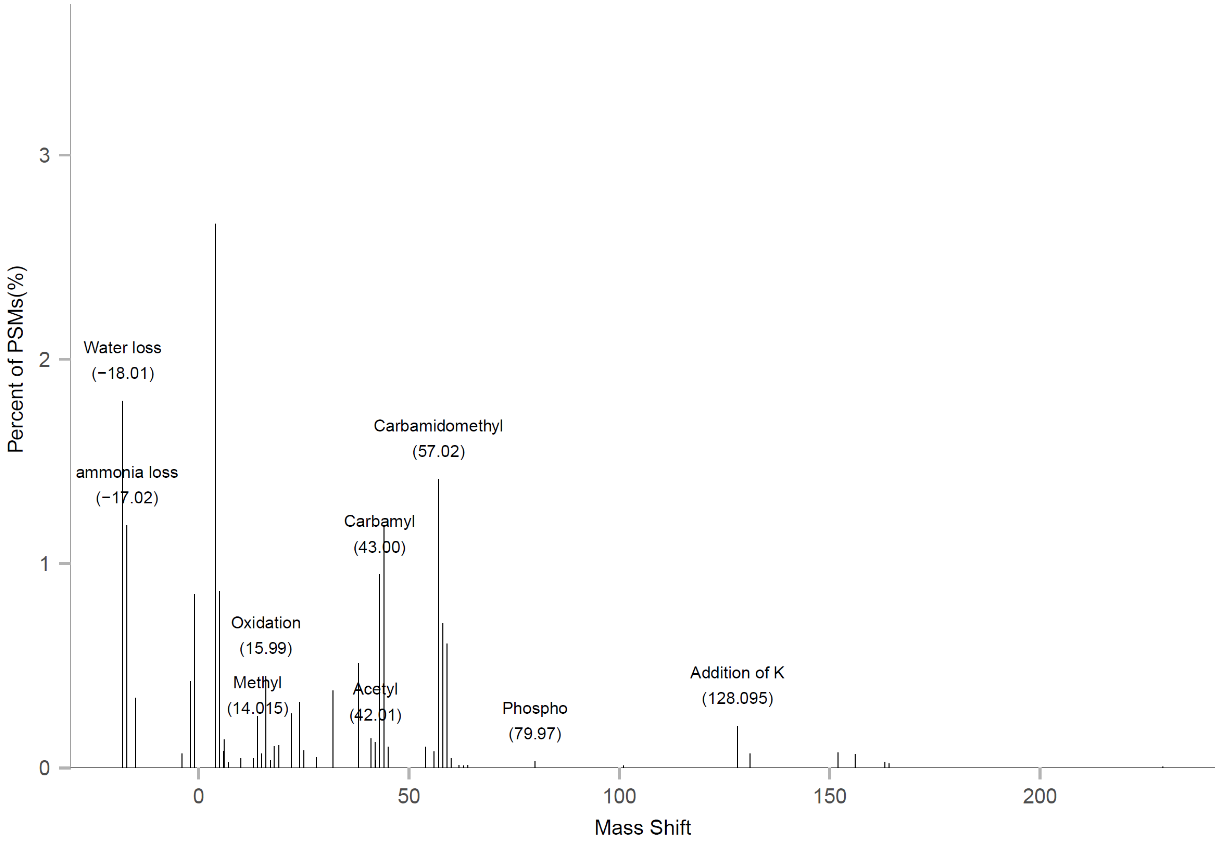


**Supplementary Figure 2.** Modifications identified using the open search workflow in FragPipe using pseudo-MS/MS spectra generated by diaTracer.

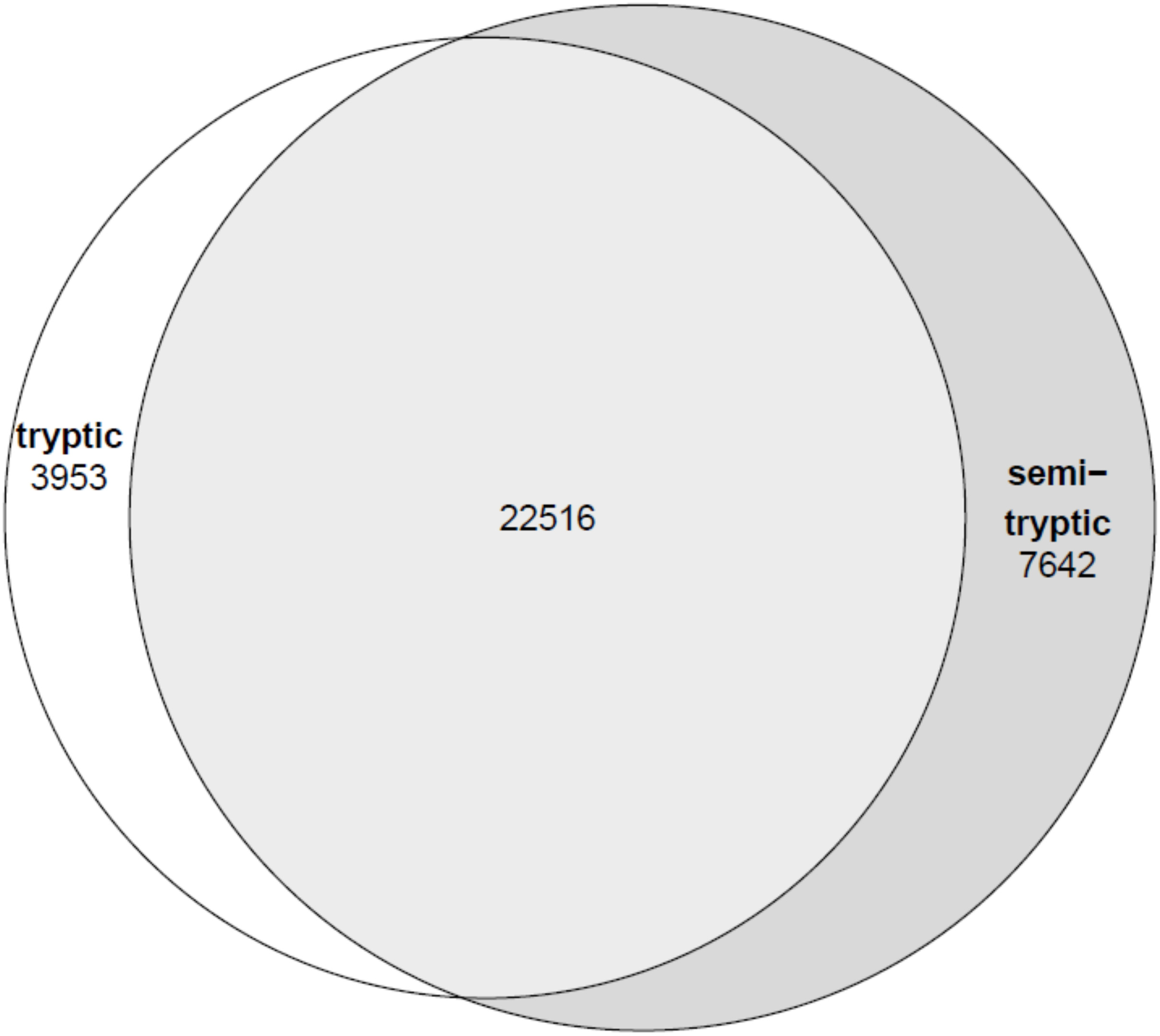


**Supplementary Figure 3.** Comparison of identified peptides between tryptic search and semi-tryptic search.

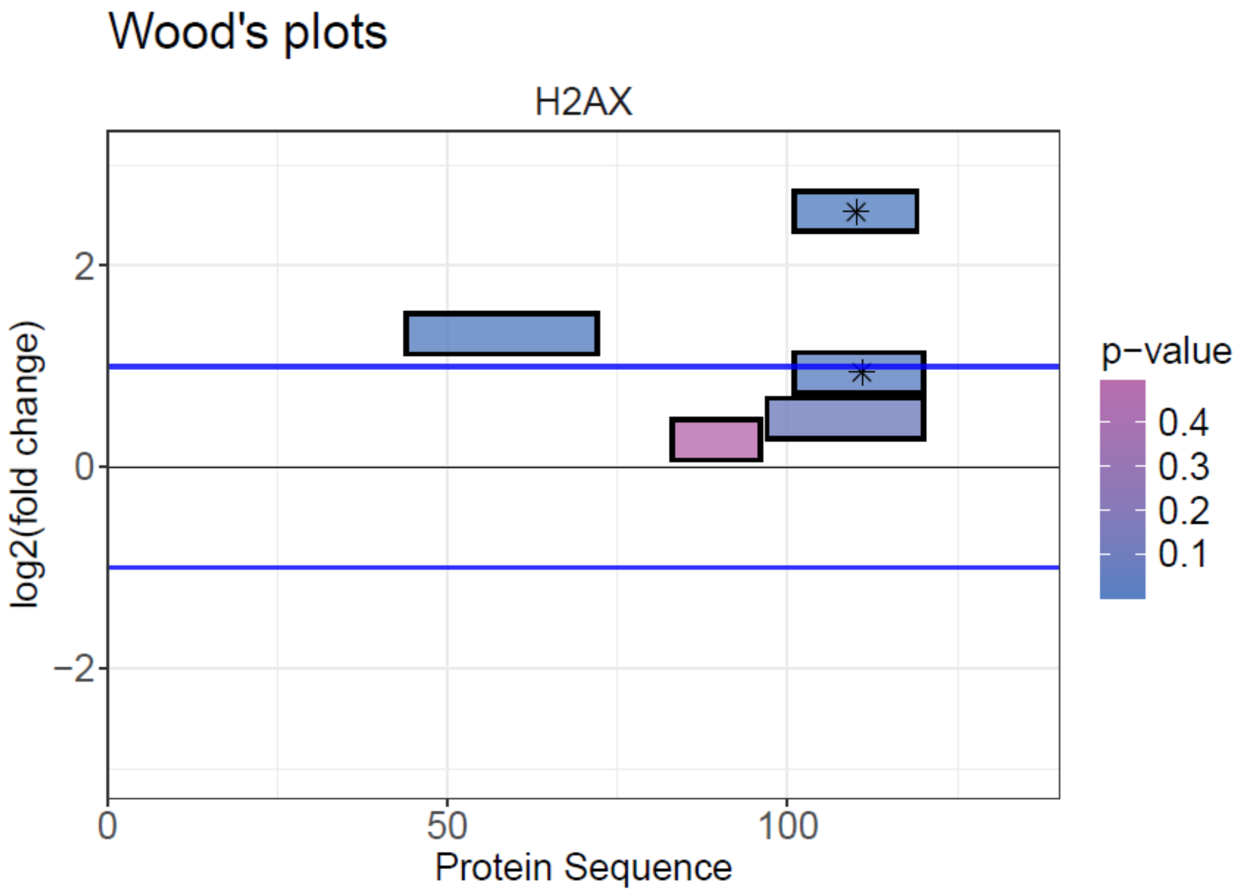


**Supplementary Figure 4.** Wood's plot of protein H2AX showing quantified tryptic and semi-tryptic (with star) peptides.

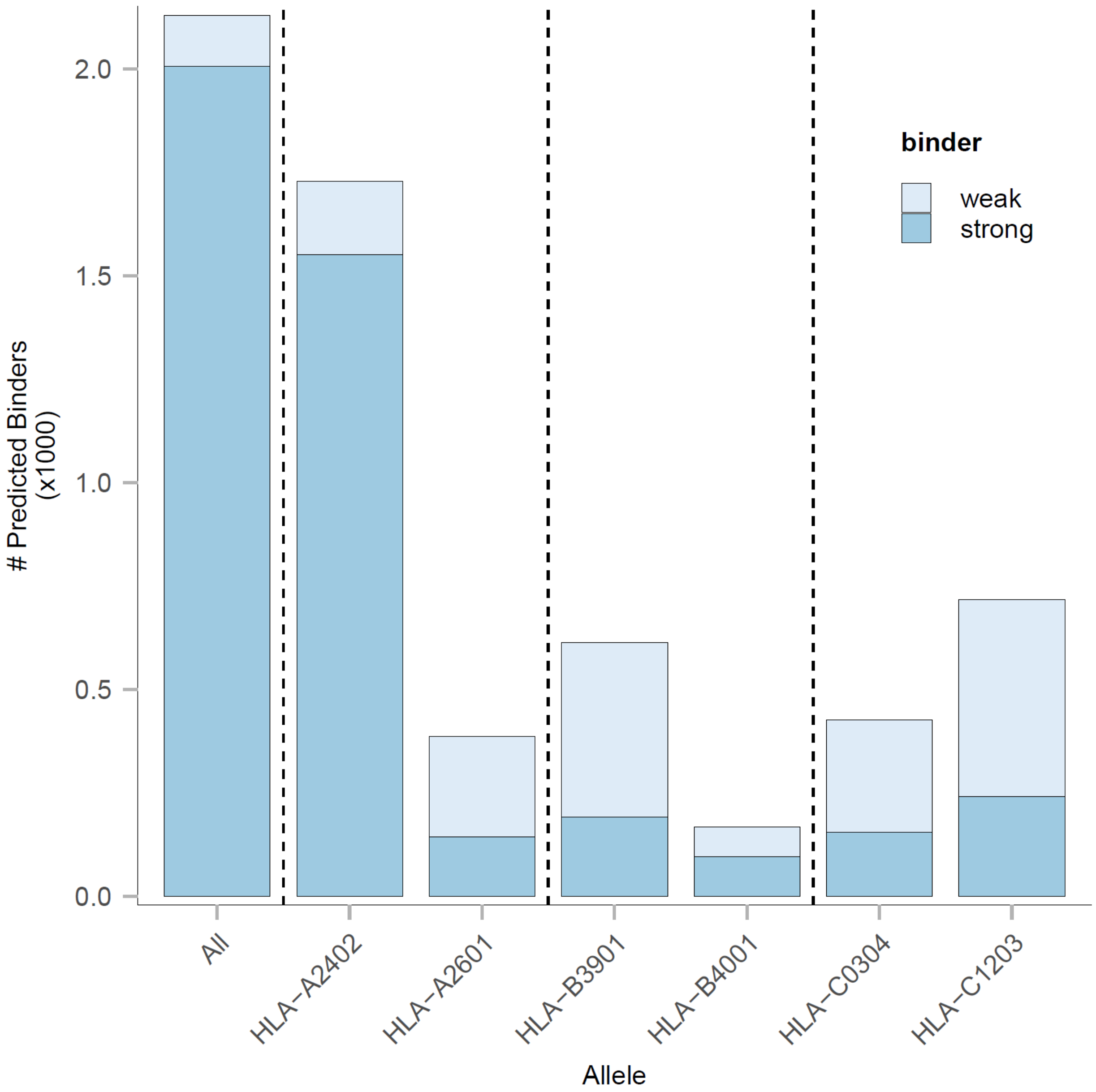


**Supplementary Figure 5.** Histogram of predicted binders of Spectronaut directDIA result from the Wahle et al. study for all HLA alleles of the corresponding sample donor, colored by binder type (light: weak binder; dark: strong binder).

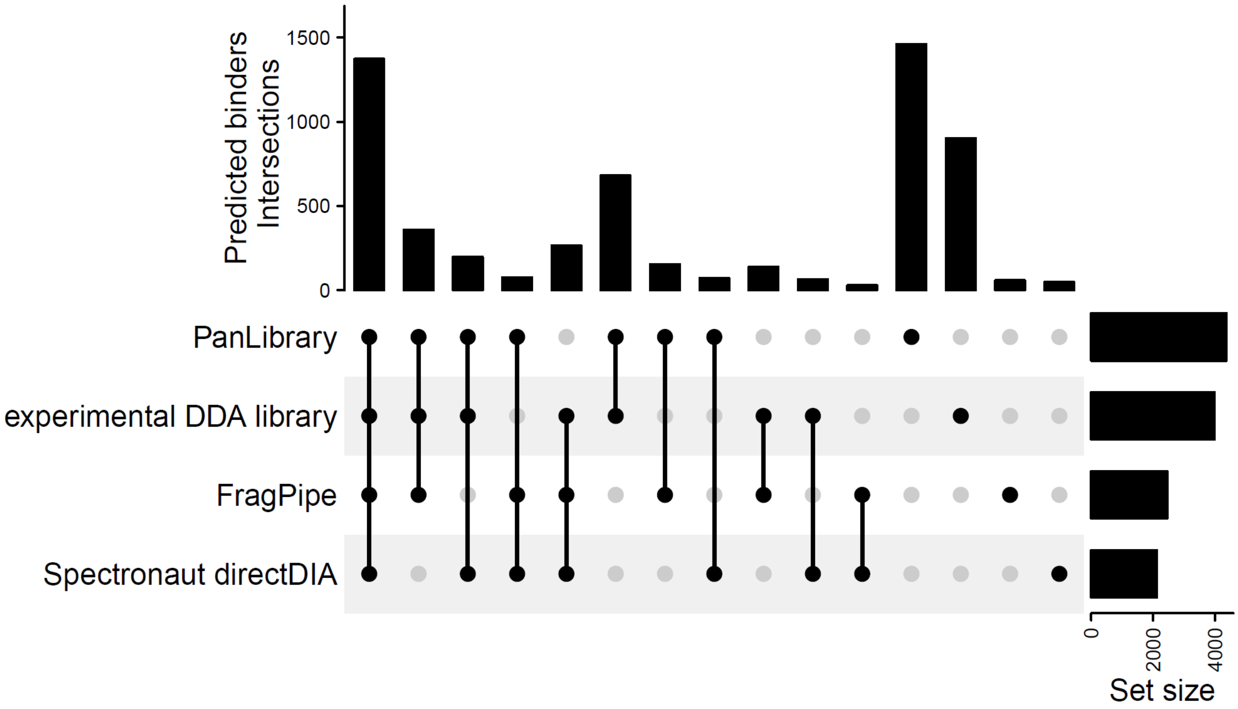


**Supplementary Figure 6.** Predicted binders overlapped with the Wahle et al. study, including the Spectronaut directDIA, experimental DDA library, and panlibrary based results.

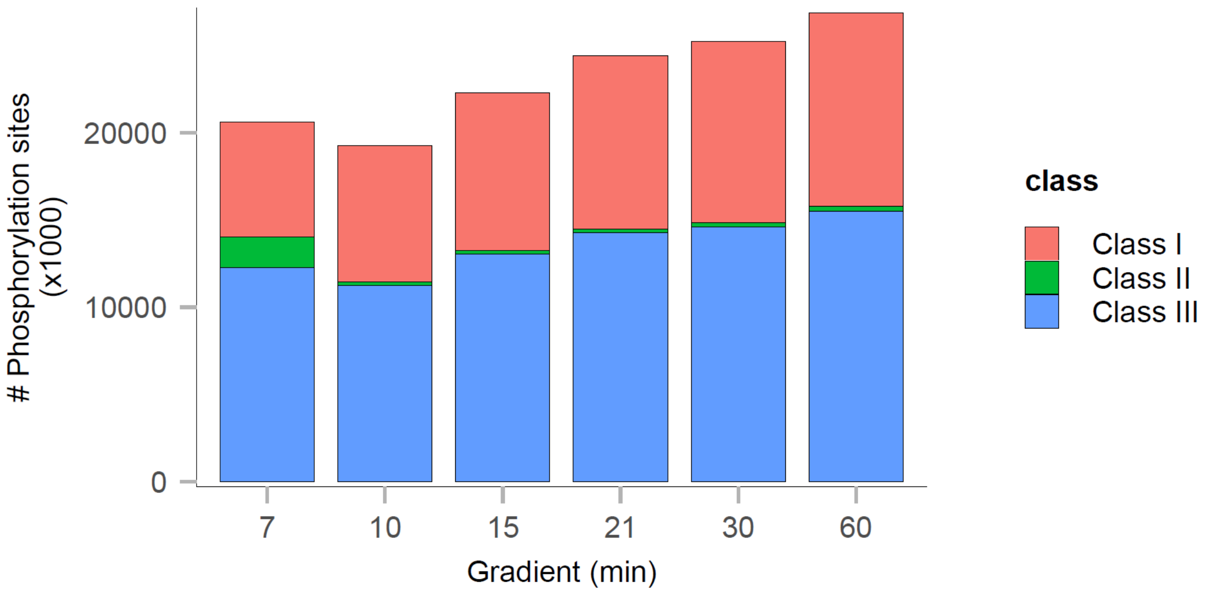


**Supplementary Figure 7.** Results of the phosphoproteome dataset. Total unique phosphorylation sites identified by FragPipe (pink: class I sites, localization probability > 0.75; green: class II sites, localization probability > 0.5 and <=0.75; blue: class III sites, localization probability <= 0.5).

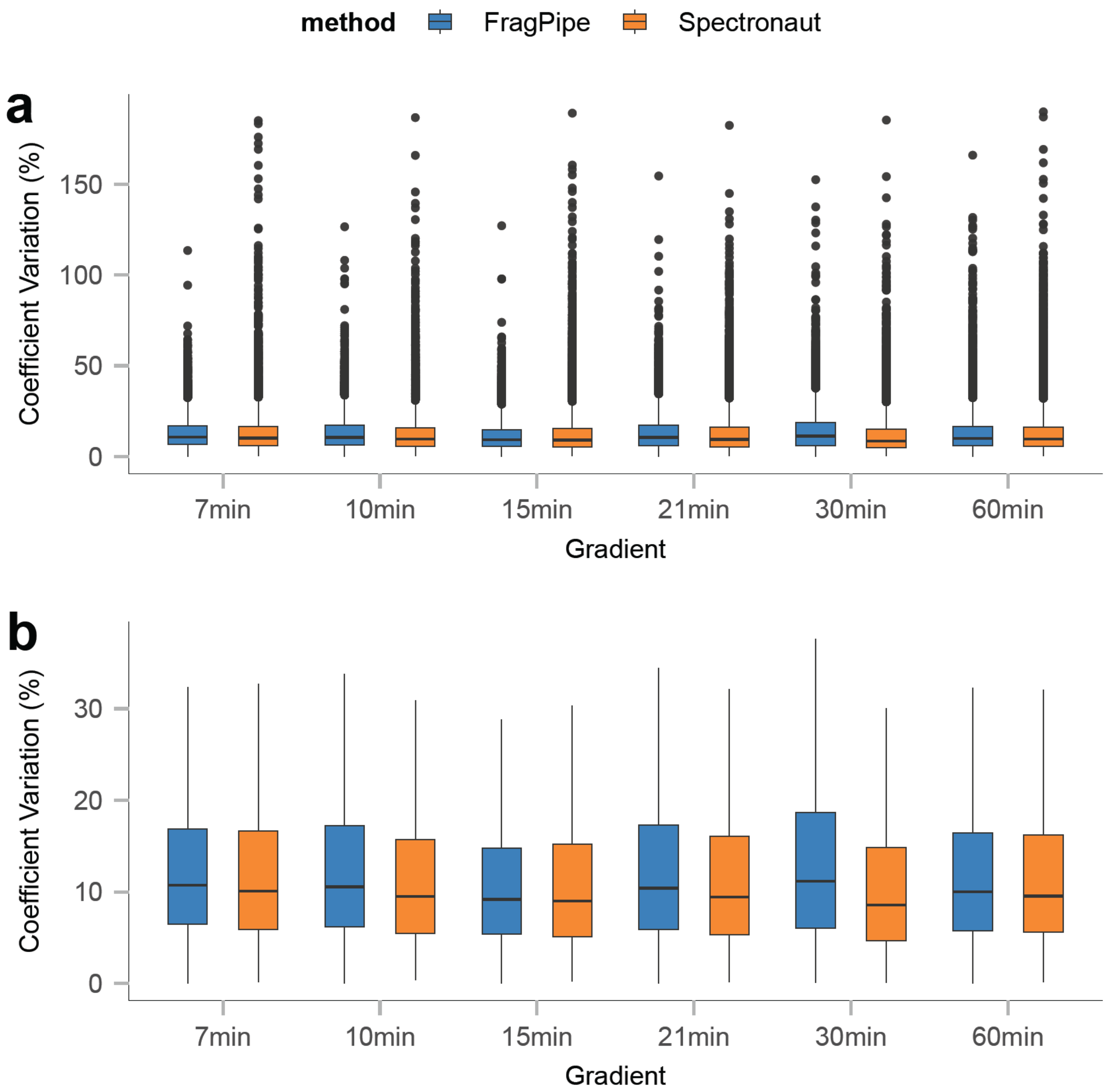


**Supplementary Figure 8.** Box plots showing coefficient variation (CV) based on phosphorylated precursor (phosphorylation probability >= 0.75) from 7-60min gradients, with colors representing processing method (blue: FragPipe with diaTracer; orange: Spectronaut). **a)** Boxplot of full range of CVs with outliers; **b)** Boxplot of CVs without outliers. The lower and upper edges of the box represent the first (Q1) and third quartiles (Q3). The interquartile range (IQR) is the box between Q1 and Q3. Data points outside this range are considered outliers and are shown as individual dots.

**a**

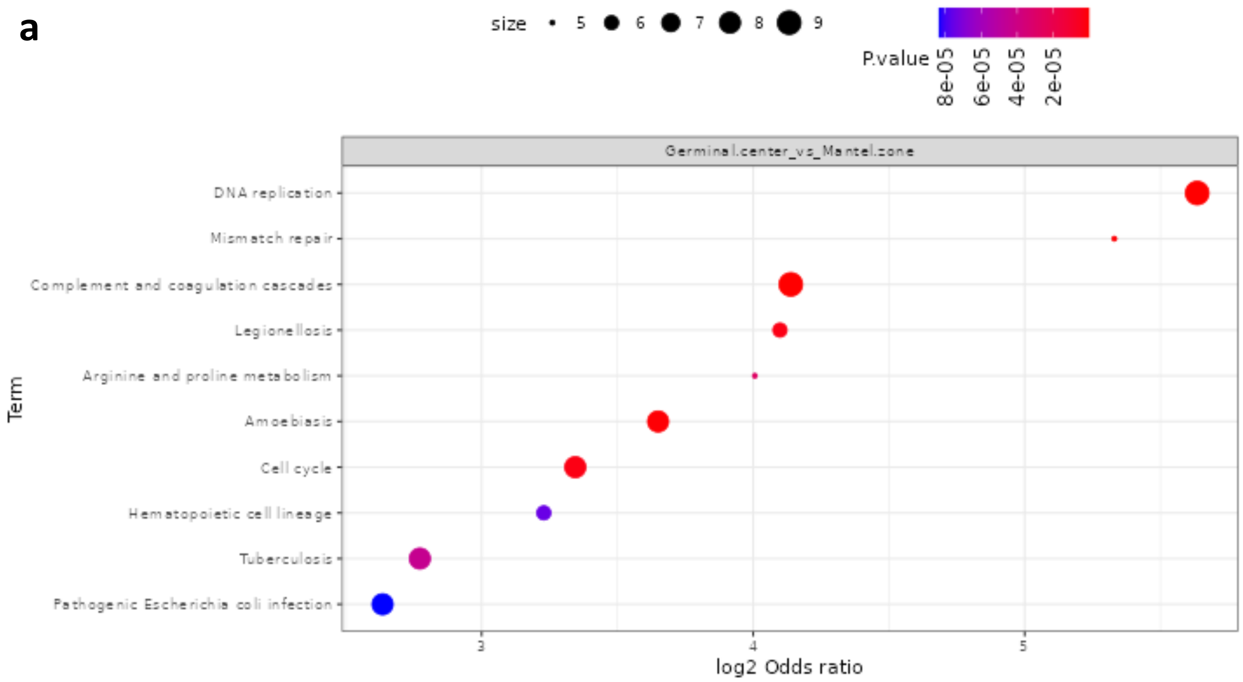
size 5 6 7 8 9
P.value 8e-05 6e-05 4e-05 2e-05
Germinal.center_vs_Mantel.zone
DNA replication
Mismatch repair
Complement and coagulation cascades
Legionellosis
Arginine and proline metabolism
Amoebiasis
Cell cycle
Hematopoietic cell lineage
Tuberculosis
Pathogenic Escherichia coli infection
Term
3 4 5
log2 Odds ratio

**b**

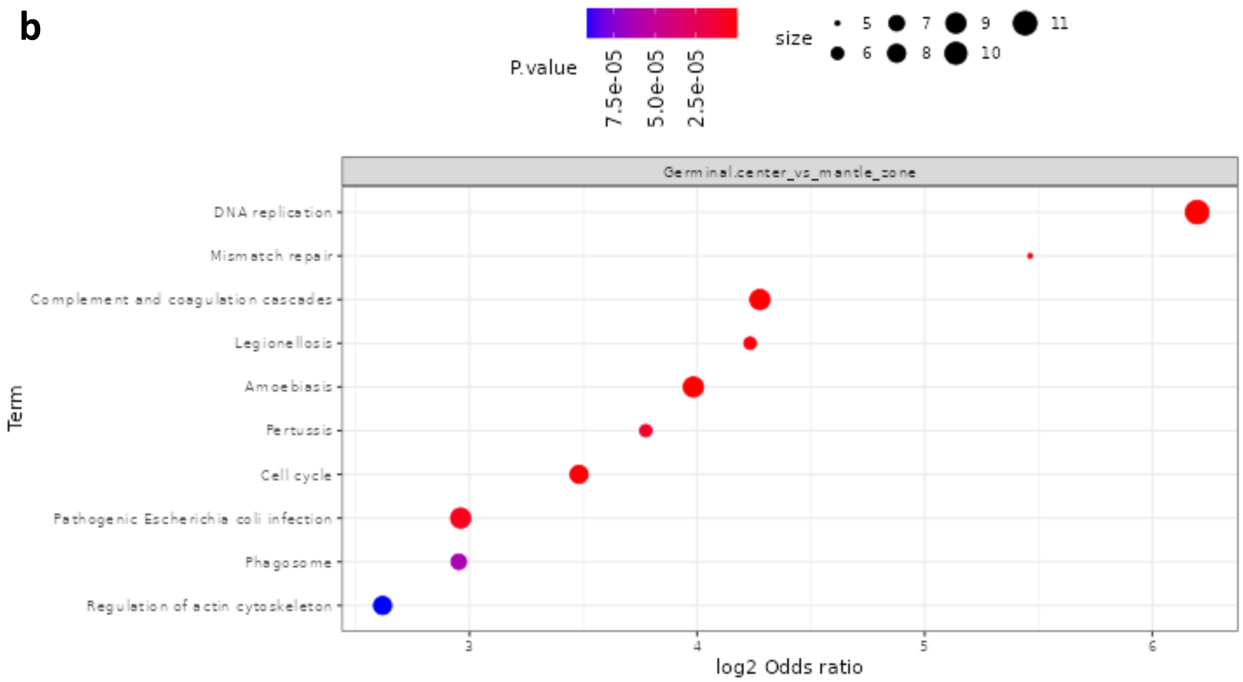
P.value 7.5e-05 5.0e-05 2.5e-05
size 5 6 7 8 9 10 11
Germinal.center_vs_mantle_zone
DNA replication
Mismatch repair
Complement and coagulation cascades
Legionellosis
Amoebiasis
Pertussis
Cell cycle
Pathogenic Escherichia coli infection
Phagosome
Regulation of actin cytoskeleton
Term
3 4 5 6
log2 Odds ratio

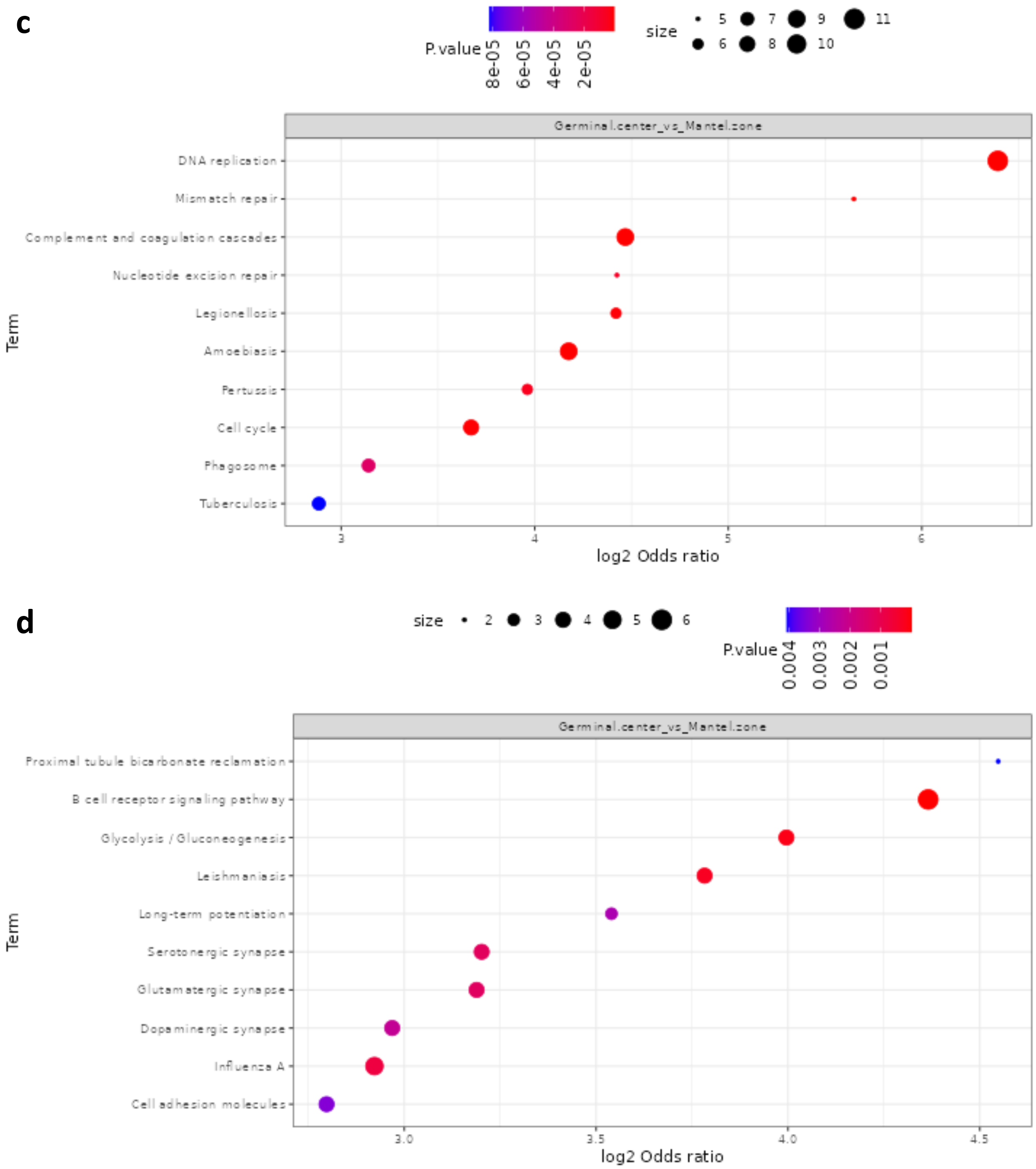
c
P.value
8e-05
6e-05
4e-05
2e-05
size
5
6
7
8
9
10
11
Germinal.center_vs_Mantel.zone
DNA replication
Mismatch repair
Complement and coagulation cascades
Nucleotide excision repair
Legionellosis
Amoebiasis
Pertussis
Cell cycle
Phagosome
Tuberculosis
Term
3
4
5
6
log2 Odds ratio
d
size
2
3
4
5
6
P.value
0.004
0.003
0.002
0.001
Germinal.center_vs_Mantel.zone
Proximal tubule bicarbonate reclamation
B cell receptor signaling pathway
Glycolysis / Gluconeogenesis
Leishmaniasis
Long-term potentiation
Serotonergic synapse
Glutamatergic synapse
Dopaminergic synapse
Influenza A
Cell adhesion molecules
Term
3.0
3.5
4.0
4.5
log2 Odds ratio

**e**

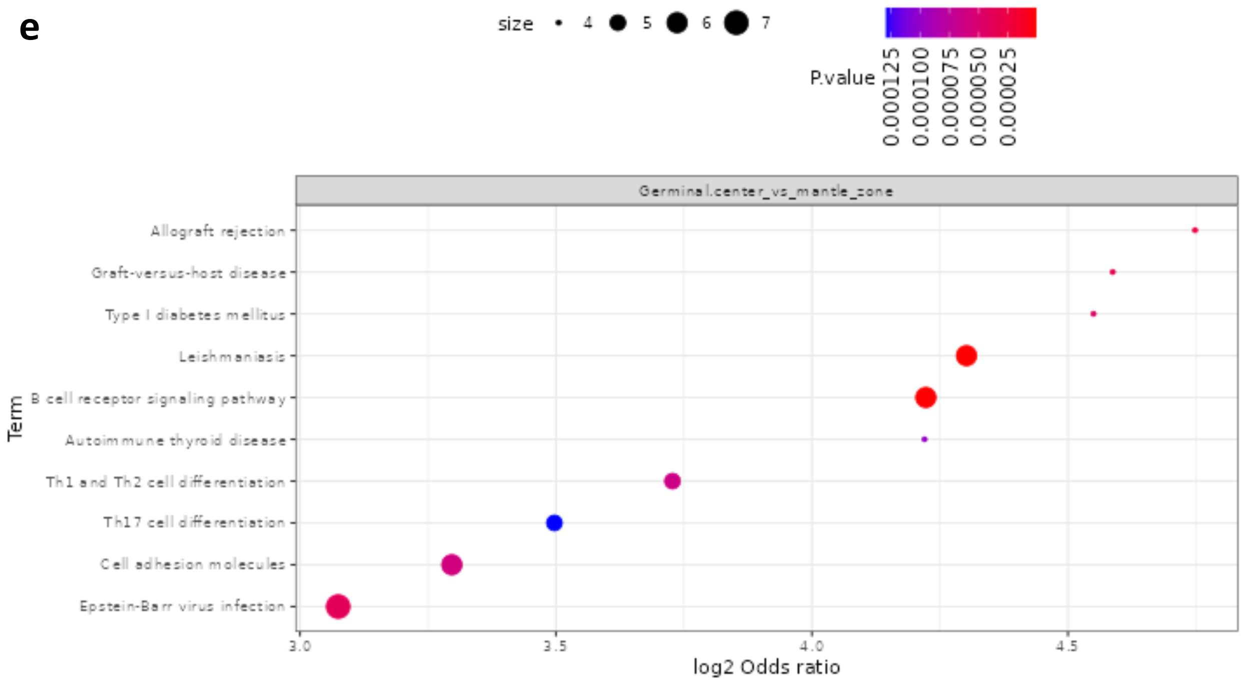


**f**

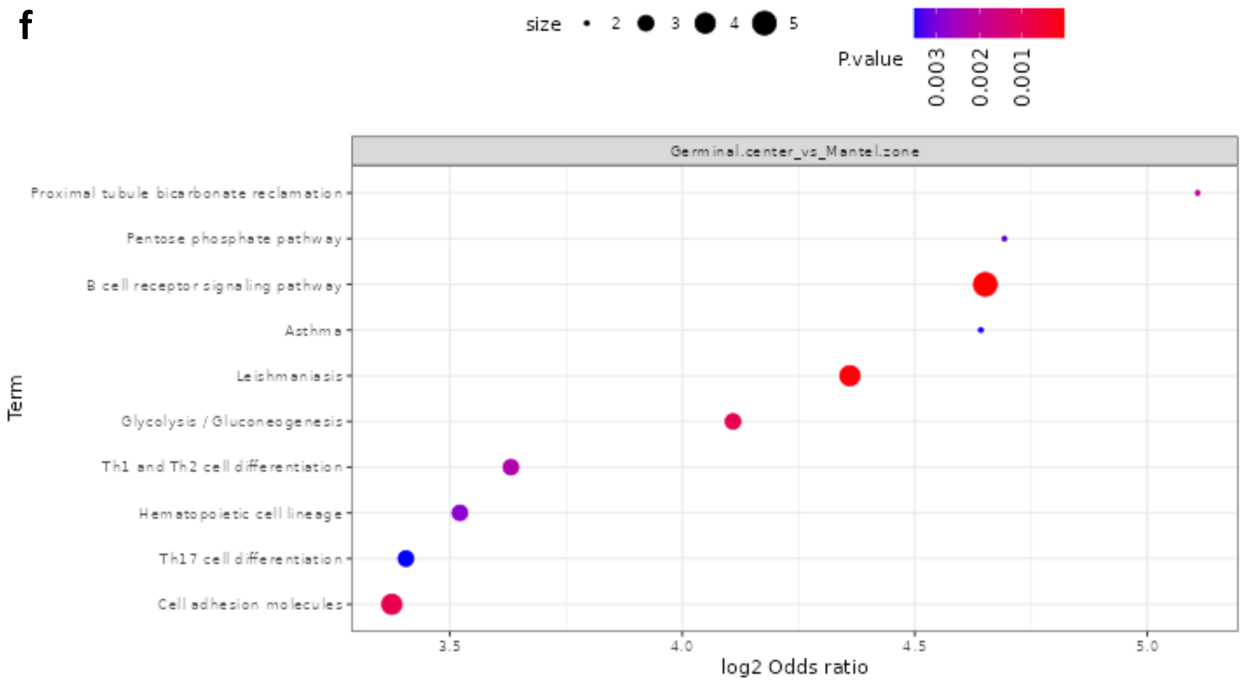


**Supplementary Figure 9.** Pathway enrichment analysis based on KEGG database of the low-input dataset between GC (germinal center) and MZ (mantel zone) groups. **a)**, **b)**, and **c)** show higher enrichment in the GC group using result from Anuar et.al, FP-diaTracer high-input library, and FP-diaTracer respectively. **d)**, **e)**, and **f)** show higher enrichment in the MZ group using result from Anuar et.al, FP-diaTracer high-input library, and FP-diaTracer respectively.
